# Large scale genomic analysis of 3067 SARS-CoV-2 genomes reveals a clonal geo-distribution and a rich genetic variations of hotspots mutations

**DOI:** 10.1101/2020.05.03.074567

**Authors:** Meriem Laamarti, Tarek Alouane, Souad Kartti, M.W. Chemao-Elfihri, Mohammed Hakmi, Abdelomunim Essabbar, Mohamed Laamarti, Haitam Hlali, Loubna Allam, Naima El Hafidi, Rachid El Jaoudi, Imane Allali, Nabila Marchoudi, Jamal Fekkak, Houda Benrahma, Chakib Nejjari, Saaid Amzazi, Lahcen Belyamani, Azeddine Ibrahimi

## Abstract

In late December 2019, an emerging viral infection COVID-19 was identified in Wuhan, China, and became a global pandemic. Characterization of the genetic variants of SARS-CoV-2 is crucial in following and evaluating it spread across countries. In this study, we collected and analyzed 3,067 SARS-CoV-2 genomes isolated from 55 countries during the first three months after the onset of this virus. Using comparative genomics analysis, we traced the profiles of the whole-genome mutations and compared the frequency of each mutation in the studied population. The accumulation of mutations during the epidemic period with their geographic locations was also monitored. The results showed 782 variant sites, of which 512 (65.47%) had a non-synonymous effect. Frequencies of mutated alleles revealed the presence of 38 recurrent non-synonymous mutations, including ten hotspot mutations with a prevalence higher than 0.10 in this population and distributed in six SARS-CoV-2 genes. The distribution of these recurrent mutations on the world map revealed certain genotypes specific to the geographic location. We also found co-occurring mutations resulting in the presence of several haplotypes. Moreover, evolution over time has shown a mechanism of mutation co-accumulation which might affect the severity and spread of the SARS-CoV-2.

On the other hand, analysis of the selective pressure revealed the presence of negatively selected residues that could be taken into considerations as therapeutic targets

We have also created an inclusive unified database (http://genoma.ma/covid-19/) that lists all of the genetic variants of the SARS-CoV-2 genomes found in this study with phylogeographic analysis around the world.

## Introduction

The recent emergence of the novel, human pathogen Severe Acute Respiratory Syndrome Coronavirus 2 (SARS-CoV-2) in China with its rapid international spread poses a global health emergency. On March 11, 2020, the World Health Organization (WHO) publicly announced the SARS-CoV-2 epidemic as a global pandemic. As of March 23, 2020, the COVID-19 pandemic had affected more than 190 countries and territories, with more than 464,142 confirmed cases and 21,100 deaths (1).

The new SARS-CoV-2 coronavirus is an enveloped positive-sense single-stranded RNA virus belonging to a large family named coronavirus which have been classified under three groups two of them are responsible for infections in mammals (2), such us: bat SARS-CoV-like; Middle East respiratory syndrome coronavirus (MERS-CoV). Many recent studies have suggested that SARS-CoV-2 was diverged from bat SARS-CoV-like (3-4). The size of the SARS-CoV2 genome is approximately 30 kb and its genomic structure has followed the characteristics of known genes of Coronavirus; the polyprotein orf1ab also known as the polyprotein replicase covers more than 2 thirds of the total genome size and structural proteins, including spike protein, membrane protein, envelope protein and nucleocapsid protein. In addition ere are also six ORFs (ORF3a, ORF6, ORF7a, ORF7b, ORF8 and ORF10) are predicted as hypothetical proteins with no associated function (5). Characterization of viral mutations can provide valuable information for assessing the mechanisms linked to pathogenesis, immune evasion and viral drug resistance. In addition, viral mutation studies can be crucial for the design of new vaccines, antiviral drugs and diagnostic tests. A previous study (6) based on an analysis of 103 genomes of SARS-CoV-2 indicates that this virus has evolved into two main types. Type L being more widespread than type S, and type S representing the ancestral version. In addition, another study (7) conducted on 32 genomes of strains sampled from China, Thailand and the United States between December 24, 2019 and January 23, 2020 suggested increasing tree-like signals from 0 to 8.2%, 18.2% and 25, 4% over time, which may indicate an increase in the genetic diversity of SARS-CoV-2 in human hosts.

Therefore, the analysis of mutations and monitoring of the evolutionary capacity of SARS-CoV-2 over time-based on a large population is necessary. In this study, we characterized the genetic variants in 3067 SARS-CoV-2 genomes for a detailed understanding of their genetic diversity and to monitor the accumulation of mutations over time with particular focus on the geographic distribution of recurrent mutations. On the other hand, we established selective pressure analysis to predict negatively selected residues which could be useful for the design of therapeutic targets. We have also created a database to share, exploit and research knowledge of genetic variants to facilitate comparison for the COVID-19 scientific community.

## Materials and Methods

### Data collection and Variant calling analysis

3067 sequences of SARS-CoV-2 were collected from the GISAID EpiCovTM (update: 02-04-2020) and NCBI (update: 20-03-2020) databases. Only complete genomes were used in this study (**Additional file 1: Table S1**). Genomes were mapped to the reference sequence Wuhan-Hu-1/2019 (NC_045512) using Minimap v2.12-r847-dirty (8). The BAM files were sorted by SAMtools sort (9), then used to call the genetic variants in variant call format (VCF) by SAMtools mpileup (9) and bcftools v1.8 (9). The final call set of the 3067 genomes, was annotated and their impact was predicted using SnpEff v 4.3t (10). First, the SnpEff databases were built locally using annotations of the reference genome NC_045512.2 obtained in GFF format from the NCBI database. Then, the SnpEff database was used to annotate SNPs and InDels with putative functional effects according to the categories defined in the SnpEff manual (http://snpeff.sourceforge.net/SnpEff_manual.html).

### Phylogentic analysis and geodistribution

The downloaded full-length genome sequences of coronaviruses isolated from different hosts from public databases were subjected to multiple sequence alignments using Muscle v 3.8 (11). Maximum-likelihood phylogenetic trees with 1000 bootstrap replicates were constructed using RaxML v 8.2.12 (39)). Heatmap for correlation analysis was performed on countries and hotspots mutations using CustVis with default settings rows scaling = variance scaling, PCA method = SVD with imputation, clustering distance for rows = correlation clustering, the method for rows = average, tree ordering for rows = tightest cluster first (12).

### Selective pressure and modelling

We used Hyphy v2.5.8 (13) to estimate synonymous and non-synonymous ratio dN / dS (ω). Two datasets of 191 and 433 for orf1ab and genes respectively were retrieved from Genbank (http://www.ncbi.nlm.nih.gov/genbank/). After deletion of duplicated and cleaning the sequences, only 91 and 39 for orf1ab and spike proteins, respectively, were used for the analysis (**Additional file 1: Table S2**). The selected nucleotide sequences of each dataset were aligned using Clustalw codon-by-codon and the phylogenetic tree was obtained using ML (maximum likelihood) available in MEGA X (14). For this analysis, four Hyphy’s methods were used to study site-specific selection: SLAC (Single-Likelihood Ancestor Counting (15), FEL (Fixed Effects Likelihood) (15), FUBAR (Fast, Unconstrained Bayesian AppRoximation) (16) and MEME (Mixed Effects Model of Evolution) (17). For all the methods, values supplied by default were used for statistical confirmation and the overall ω value was calculated according to ML trees under General time reversible model (GTR model). The CI-TASSER generated models (https://zhanglab.ccmb.med.umich.edu/COVID-19/) of nonstructural proteins (nsp3, nsp4, nsp6, nsp12, nsp13, nsp14 and nsp16 of orf1ab were used to highlight the sites under selective pressure on the protein. On the other hand, the cryo-EM structure with PDB id 6VSB was used as a model for the spike protein in its prefusion conformation. Structure visualization and image rendering were performed in PyMOL 2.3 (Schrodinger LLC).

### Pangenome construction

115 proteomes of the genus *Betacorononavirus* were obtained from the NCBI database (update: 20-03-2020), of which 83 genomes belonged to the SARS-CoV-2 species and the rest distributed to other species of the same genus publicly available (**Additional file 2: Table S3**). These proteomes were used for the construction of pangenome at the inter-specific scale of *Betacoronavirus* and intra-genomic of SARS-CoV-2. The strategy of best reciprocal BLAST results (18) was implemented to identify all of the orthologous genes using Proteinortho v6.0b (19). Proteins with an identity above 60% and sequence coverage above 75% with an e-value threshold below 1e-5 were used to be considered as significant hits.

## Results

### SARS-CoV-2 genomes used in this study

In this study, we used 3,067 SARS-CoV2 complete genomes collected from GISAID EpiCovTM (update: 02-04-2020) and NCBI (update: 20-03-2020) databases. These strains were isolated from 55 countries (**Fig 1A**). The most represented origin was American strains with 783 (25.53%), followed by strains from England, Iceland, and China with 407 (13.27%), 343 (11.18%), 329 (10.73%), respectively. The date of isolation was during the first three months after the appearance of the SARS-CoV-2 virus, from December 24, 2019, to March 25, 2020 (**Fig 1B**). Likewise, about two-thirds of these strains collected in this work were isolated during March.

**Figure 1:**
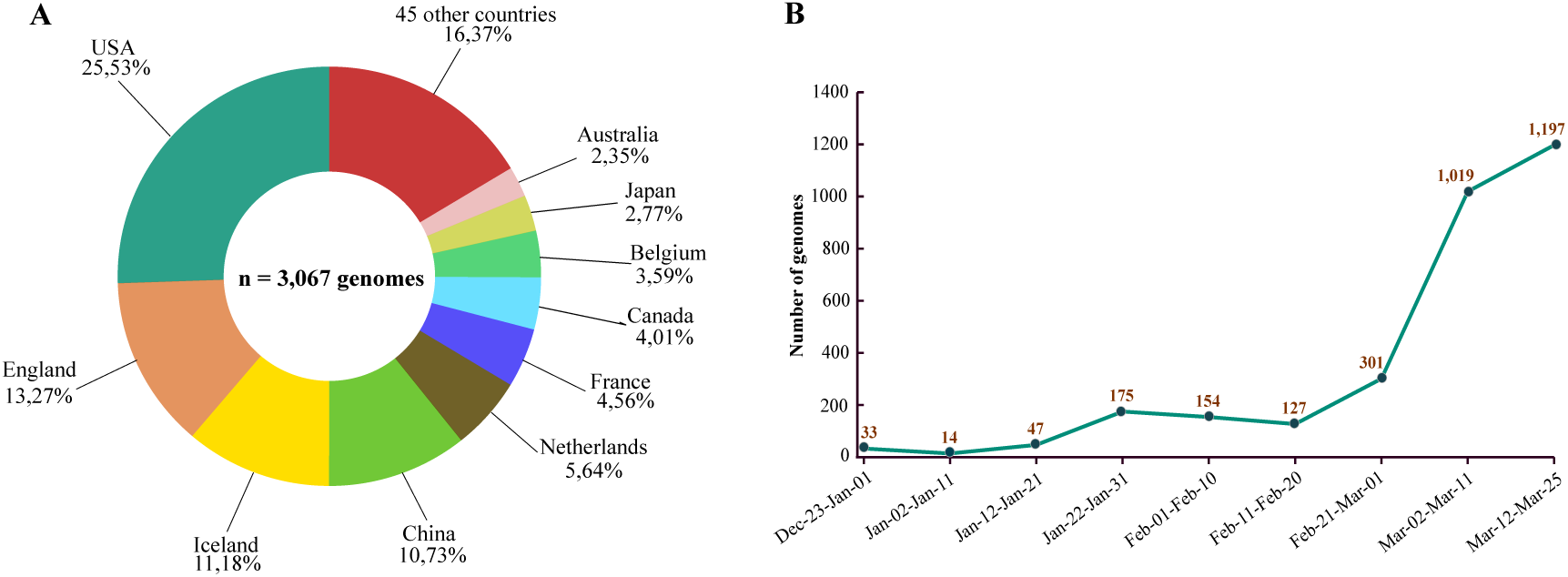
Distribution of the genomes of the 3,067 genomes used in this study by county and date of isolation. A) The pie chart represents the percentage of genomes used in this study according to their geographic origins. The colors indicate different countries. B) Number of genomes of complete pathogens, distributed over a period of 3 months from the end of December to the end of March

### Allele frequencies revealed a diversity of genetic variants in six SARS-Cov-2 genes

To study and follow the appearance and accumulation of mutations, we have traced the profiles of these mutations and compared their frequencies in the population studied. Remarkably, compared to the Wuhan-Hu-1/2019 reference sequence, a total of 782 variant sites were identified, including 512 (65.47%) non-synonymous mutations, 222 (28.38%) synonymous mutations, and four (0.51%) deletion mutation effect. The rest (5.64%) distributed to the intergenic regions. Frequency analysis of the mutated alleles revealed the presence of 68 recurrent mutations with a prevalence greater than 0.006 (0.06% of the population), which corresponds to at least 20 / 3,067 genomes of SARS-CoV-2. Focusing on recurrent non-synonymous mutations, 38 was found and distributed in six genes with variable frequencies (**Fig 2**), of which the gene coding for replicase polyprotein (orf1ab), spike protein, membrane glycoprotein, nucleocapsid phosphoprotein, ORF3a, and ORF8. Overall, orf1ab harbored more non-synonymous mutations compared to the other five genes with 22 mutations, including three mutations located in nsp12-RNA-dependent RNA polymerase (RdRp) (M4555T, T4847I and T5020I), three in nsp13-helicase (V5661A, P5703L and M5865V), two in nsp5-main proteinase (G3278S and K3353R), two in nsp15-EndoRNAse (I6525T, Ter6668W), two in nsp3-multi domains (A876T and T1246I), one in nsp14-exonuclease (S5932F) and one in nsp4-transmembrane domain 2 (F3071Y). Likewise, spike protein harbored three frequent mutations, including V483A in receptor-binding domain (RBD). The rest of the mutations were found in nucleocapsid phosphoprotein (S193I, S194L, S197L, S202N, R203K and G204R), ORF3a (S193I, S194L, S197L, S202N, R203K and G204R), membrane glycoprotein (D3G and T175M) and ORF8 (V62L and L84S).

**Figure 2:**
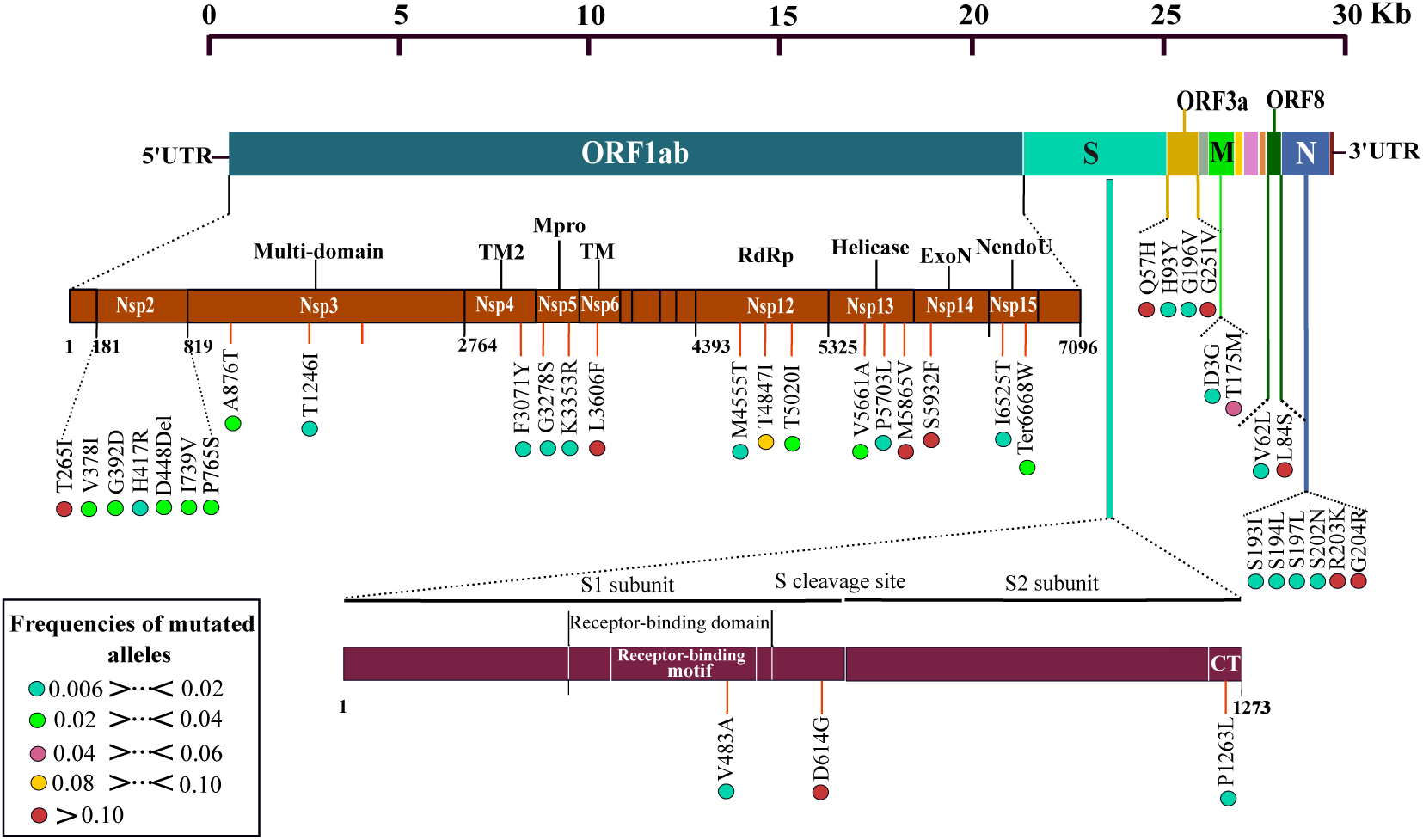
Schematic representation of the SARS-CoV-2 genome with recurrent non-synonymous mutations. The brown and garnet diagrams illustrate the non-structural proteins (nsp1 to nsp 16) of the ORF1ab protein and the two subunits of the spike (S) protein, respectively. Recurrent mutations represented by vertical lines. The frequency of each mutation in the population is presented by color coded circles.

### Identification of ten hyper-variable genomic hotspot in SARS-CoV-2 genomes

Interestingly, among all recurrent mutations, ten were found as hotspot mutations with a frequency greater than 0.10 in this study population (**Fig 2**). The most represented was D614G mutation at spike protein with 43.46% (n = 1.333) of the genomes, the second was L84S (at ORF8) found in 23.21% (n = 712). Thus, the gene coding for orf1ab had four mutations hotspots, including S5932F of nsp14-exonuclease, M5865V of nsp13 helicase L3606F of nsp6 transmembrane domain and T265I of nsp2 found with 17.02%, 16.56%, 14.38% and 10.66% of the total genomes, respectively. For the four other hotspot mutations were distributed in ORF3a (Q57H and G251V) and nucleocapsid phosphoprotein (R203K and G204R).

### Geographical distribution and origin of mutations worldwide

3067 genomes were dispersed in different countries with different genotype profiles. We performed a geo-referencing mutation analysis to identify region-specific loci. Remarkably, China and USA were the countries with the highest number of mutations 301 and 296 (38,19 % and 37,56 % of the total number of mutations) including 140 (17,76%) and 229 (29%) singleton mutations specific to China and USA genomes respectively, followed by Malaysia and France with 3,6% and 2,4%, respectively.

It is interesting to note that among the 55 countries, 21 harbored singleton mutations. **(Additional file 3: Table S4)** illustrates the detailed singleton mutations found in these countries. The majority of the genomes analyzed carried more than one mutation. However, among the recurrent non-synonymous, synonymous, deletion and intergenic mutations, we found G251V (in ORF3a), and S5932F (in ORF1ab) present on all continents except Africa (**Fig 3)**. While F924F, L4715L (in orf1ab), D614G (in spike) appeared in all strains except those from Asia. In Algeria, the genomes harbored mutations very similar to those in Europe, including two recurrent mutations T265I and Q57H of the ORF3a. Likewise, the European and Dutch genomes also shared ten recurrent mutations. On the other hand, continent-specific mutations have also been observed, for example in America, we found seven mutations shared in almost all genomes. Besides, two mutations at positions 28117 and 28144 were shared by the Asian genomes, while four different positions 1059, 14408, 23403, 25563 and 1397, 11083, 28674, 29742 were shared by African and Australian genomes (Supplementary material). The majority of these mutations are considered to be transition mutations with a high ratio of A substituted by G. The genome variability was more visible in China and USA than in the rest of the world.

**Figure 3:**
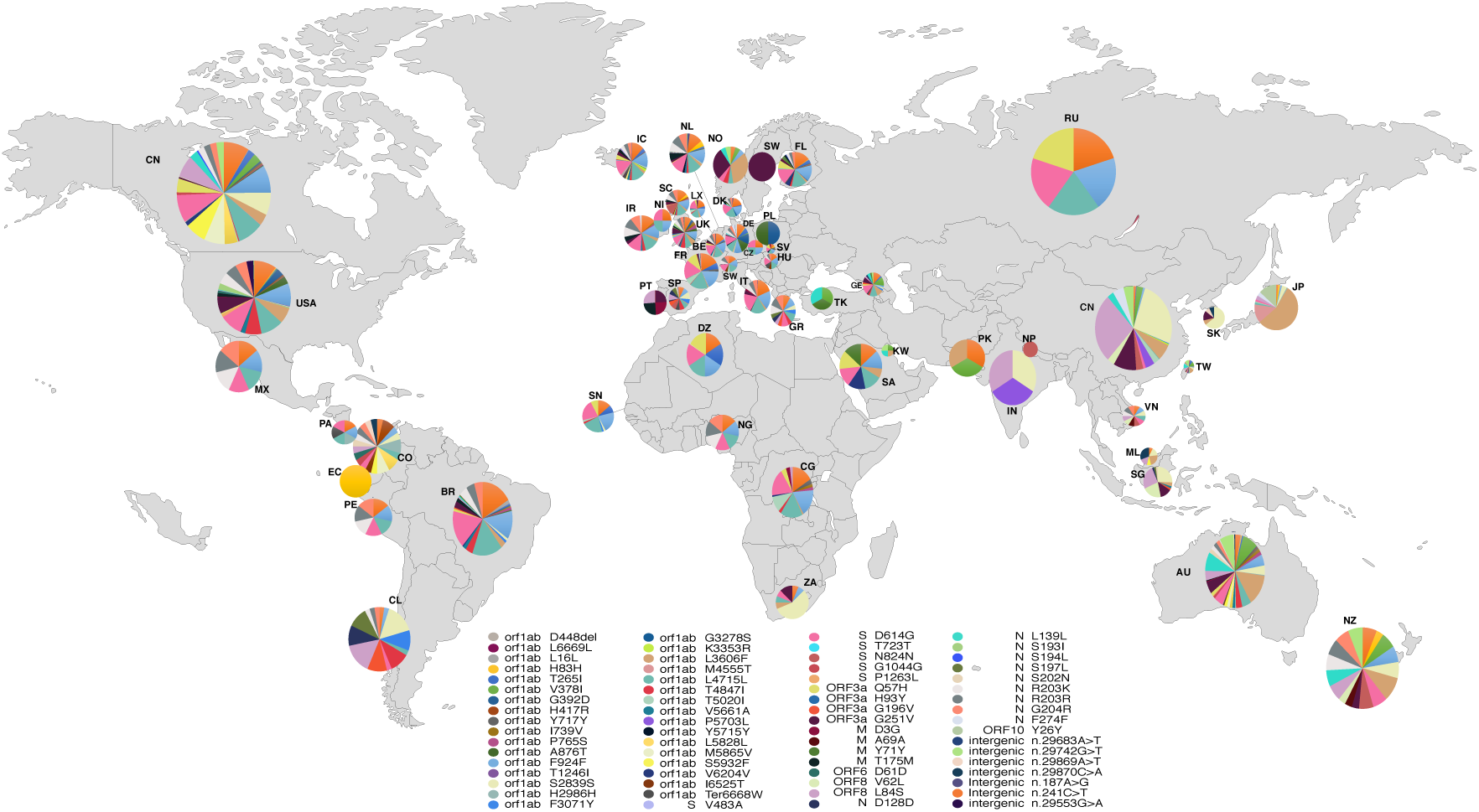
Map showing Geographical distribution of hotspot mutation in the studied population worldwide. The pie charts show the relative frequencies of haplotype for each population. The haplotypes are color coded as shown in the key. The double-digit represent countries’ two letters code. The circle’s size was randomly generated with no association with the number of genomes in each country.

SARS-CoV-2 genomes also harbored three co-occurrent mutations R203K, R203R and G204R in the N protein and were present in all continents except Africa and Asia (besides Taiwan).

### Evolution of mutations over time

We selected the genomes of the SARS-CoV-2 virus during the first three months after the emergence of this virus (December 24 to March 25). We have noticed that the mutations have accumulated at a relatively constant rate (**Fig 4**). The strains selected at the end of March showed a slight increase in the accumulation of mutations with an average of 11.34 mutations per genome, compared to the gnomes of February, December and January with an average number of mutations of 9.26, 10.59 and 10.34 respectively. The linear curve in Figure 5 suggests a continuous accumulation of SNPs in the SARS-CoV-2 genomes in the coming months. This pointed out that many countries had multiple entries for this virus that could be claimed. Thus in the deduced network demonstrated transmission routes in different countries.

**Figure 4:**
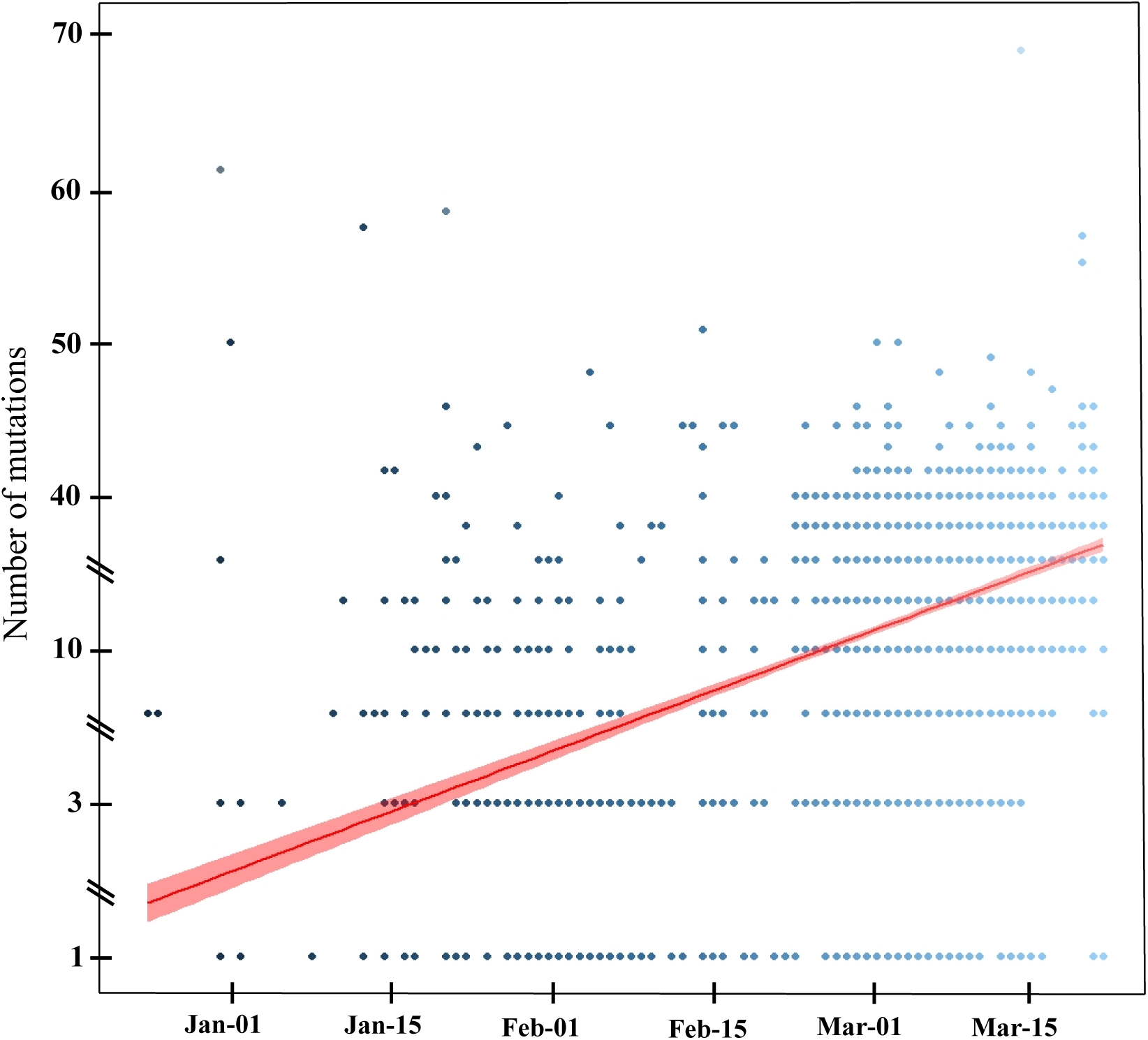
The graph represents substitutions accumulation in a three months period. The accumulation of mutations increases linearly with time. The dots represent the number of mutations in a single genome. All substitutions were included non-synonymous, synonymous, intergenic.

**Figure 5:**
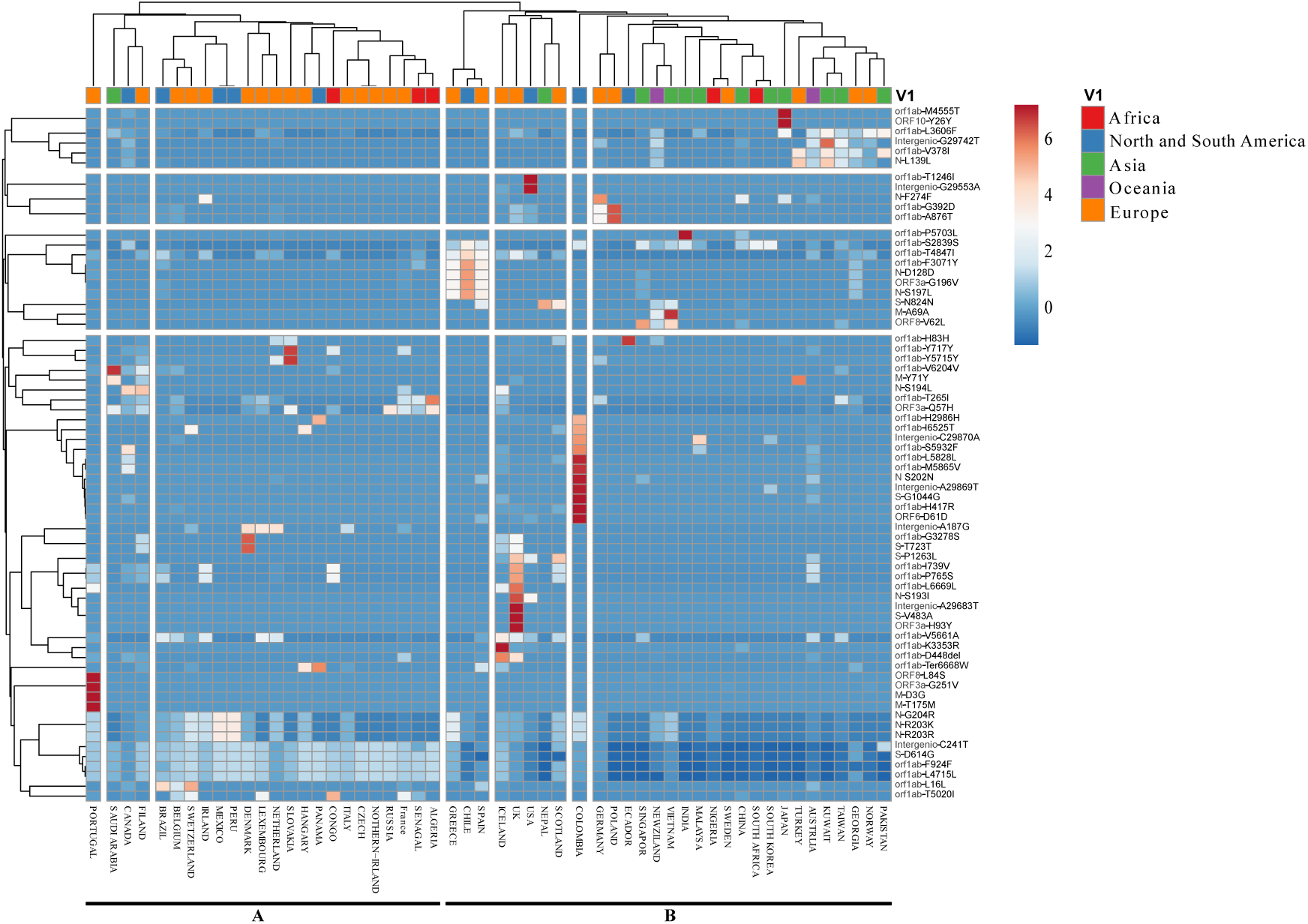
Heatmap demonstrating Correlation between mutations and geographical distribution of the analyzed genomes. The correlation was applied to a data set of 69 most recurrent mutations with different distribution in all 56 countries divided into two distinct cluster A and B. The color scale indicates the significance of correlation with blue and orange colors indicating the highest and lowest correlation. The red, yellow and orange colors in the horizontal bar represent the continent of origin.

The study of mutations accumulation over time showed a higher number of mutations in the middle of the outbreak (end of January). At the same time, an increase in the number of mutations in early April was also observed. The first mutations to appear were mainly located in the intergenic region linked to the nucleocapsid phosphoprotein and the orf8 protein. The T265I, D614G and L84S hotspot mutations located in orf1ab and Spike proteins respectively were introduced into the virus for the first time in late February.

### Phylogeographical analysis of SARS-CoV-2 genomes

The phylogenetic tree based on the whole genome alignment demonstrates that SARS-CoV-2 is wildly disseminated across distinct geographical location. The results showed that several strains are closely related even though they belong to different countries. Which indicate likely transfer events and identify routes for geographical dissemination. For phylogenetic tree (http://genoma.ma/covid-19/) showed multiple introduction dates of the virus inside the USA with the first haplotype introduced related to the second epidemic wave in China.

Using correlation analysis between most recurrent mutations and countries distribution (**Fig 5**). We observed that most recurrent mutations clusters could be divided into four groups; the bigger cluster compromised nine mutations from the ten hotspots, while the first cluster harbored only the orf1ab mutation L3606F.

Meanwhile, geo clustering by geographic location showed two distinct clusters (**Fig 5**), cluster A grouping countries from Europe with those from America and Africa. However, Asia was only represented by Saudi Arabia. Cluster B in the other hand contained the majority of countries from the Asian and Australian continents. it is also harboring a sub-cluster containing the UK, USA, and Ireland which was previously demonstrated to contain a high number of mutations.

On the other hand, mutations as V378I and L3606F (in orfab1), 29742 C>T (intergenic), L139L in (in nucleocapside) were mainly correlated with Pakistan, Norway, Georgia, Taiwan, Kuwait, Australia, and Turkey while (S2839S, F3071Y and T4847I), D128D and G196V mutations in orf1ab, nucleocapsid, ORF3a, respectively, were mainly present in Spain, Chile, and Greece. However, cluster harboring D614G (in spike), F924F (in orf1ab), and L4715L (in orf1ab) mutations, showed no correlation and were scatted through all countries especially those from Europe. A high correlation with a specific mutation was observed within Portugal, Saudi Arabia, Slovakia, Iceland, UK, USA, Colombia, Ecuador, Vietnam, Japan genomes.

### Selective pressure analysis

Selective pressure on orf1ab, gene harbored a high rate of mutations and on the Spike gene, indicated a single alignment-wide ω ratio of 0.571391 and 0.75951 for spike and or1ab, respectively. Most sites for both genes had ω <1 values, indicating purifying selection. In orf1ab, we estimated eight sites under negative selection pressure (696, 1171, 2923, 3003, 3715, 5221, 5704 and 6267) and three sites under positive selection pressure (1473, 2244 and 3090). For spike, we found seven sites under negative selection pressure (215, 474, 541, 809, 820, 921 and 1044), and only one site under negative selection pressure (**Table 1**).

**Table 1:** Selective pressure analysis on the spike and orf1ab genes of SARS-CoV-2.

| **Genes** | **$\omega$** | **FEL method** |  | **MEME method** | **SLAC method** |  | **FUBAR method** |  |
| --- | --- | --- | --- | --- | --- | --- | --- | --- |
| **Spike** | 0.571391 | **PS** | **NS** | **PS** | **PS** | **NS** | **PS** | **NS** |
|  |  | - | Codons 215, 474, 809, 820, 921, | - | - | - | Codon 5 | Codons 215, 474, 541, |
| **orflab** | 0.75951 | **PS** | **NS** | **PS** | **PS** | **NS** | **PS** | **NS** |
|  |  | Codon 2244 | Codons 1171, 2923, 3003, 3715, 5221, 5704, 6267, 6961 | Codon2244 | - | - | Codons 1473, 2244, 3090 | - |

The modelling results of orf1ab showed that the sites with positive selections were distributed in nsp3 and nsp4, while the negatively selected codons were located in nsp3, nsp4, nsp6, nsp12, nsp13, nsp14 and nsp16 (**Fig 6**). In spike, the only negatively selected residue was observed in the RBD region (**Fig 7**).

**Figure 6:**
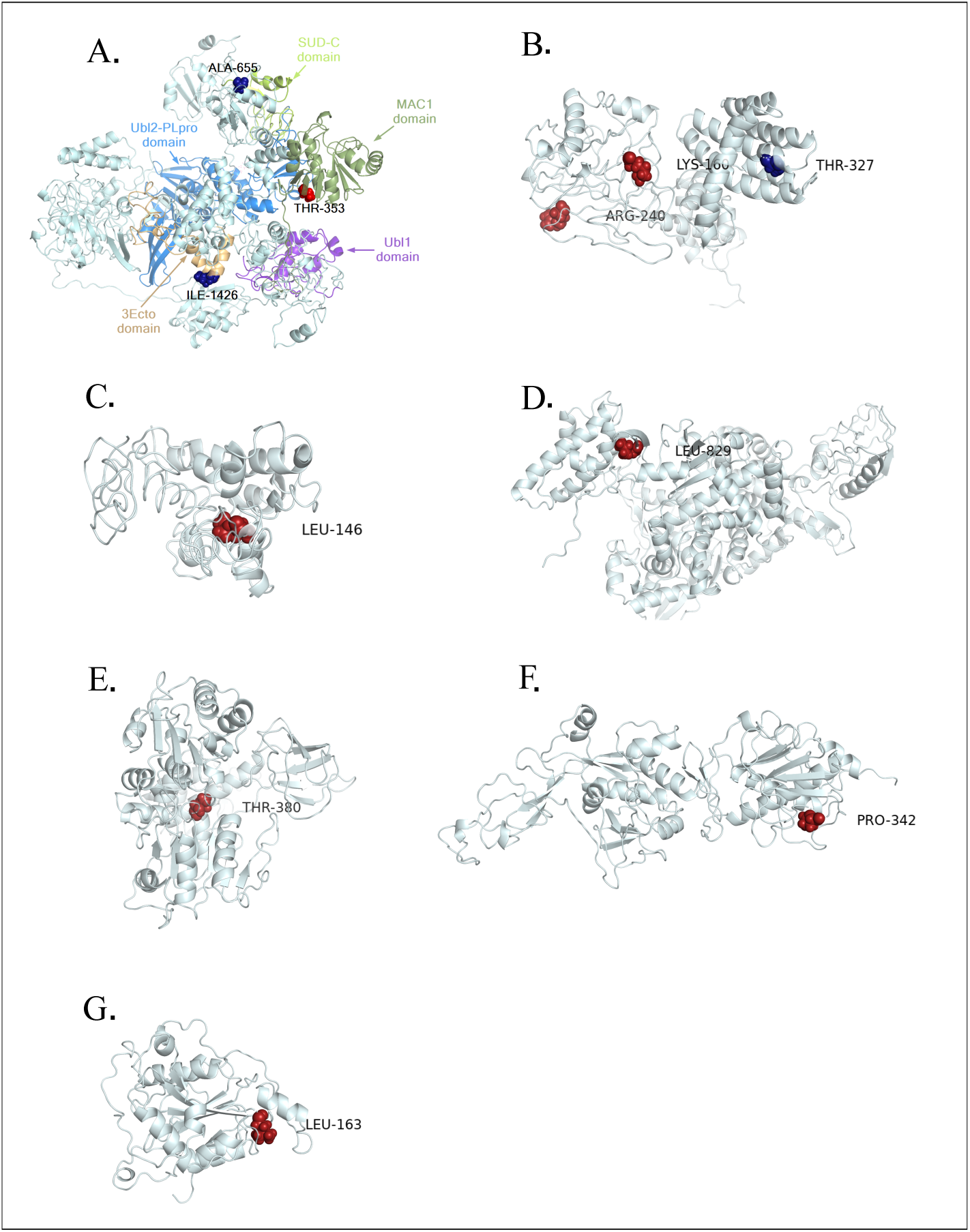
Structural view of selective pressure in orf1ab gene. The residue under the positive and negative selection is highlighted in blue and red respectively. The modeling of orf1ab non-structural proteins (NSP3, NSP4, NSP6, NSP12, NSP13, NSP14, and NSP16) harboring residues under pressure selection was produced using CI-TASSER. A. The NSP3 domains MAC1, Ubl1, Ubl2-PLpro, and SUD-C are color-coded in the 3D representation. The residues Ile-1426 and Ala-655 under negative selection are located respectively on 3Eco and SUD-C domains while Thr-353 residue under positive selection is shown on the MAC1 domain, B. 3D representation of the NSP4 protein, C. 3D representation of the NSP6 protein, D. 3D representation of the NSP12 protein, E. 3D representation of the NSP13 protein, F. 3D representation of the NSP14 protein, G. 3D representation of the NSP16 protein.

**Figure 7:**
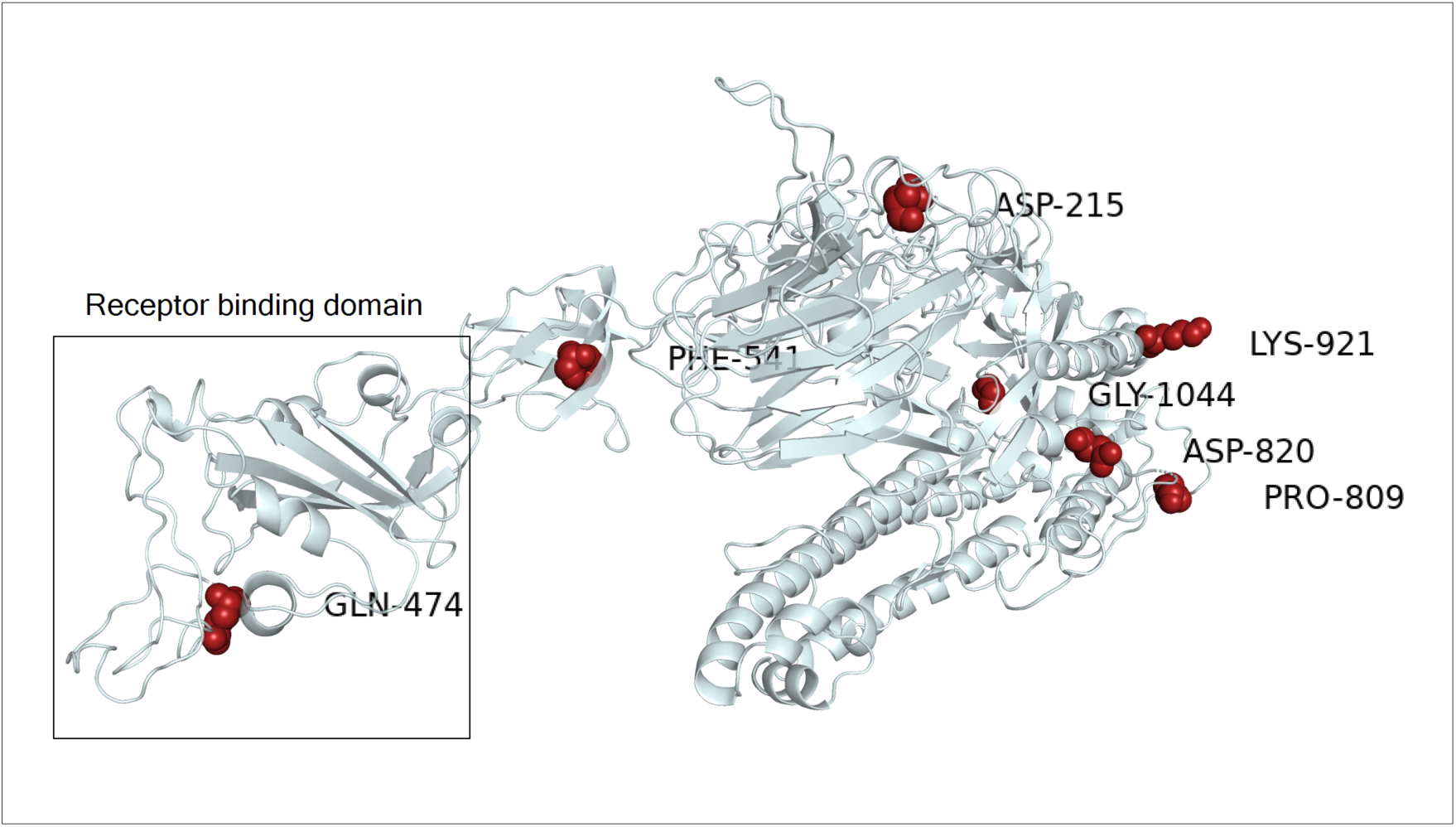
Structural view of selective pressure in spike gene. The negatively selected site in spike protein is highlighted in red. The only amino acid residue selected negatively on the receptor-binding domain corresponds to GLN-474. The cryo-EM structure with PDB id 6VSB was used as a model for the gene S in its prefusion conformation.

### Inter and intra-specific pan-genome analysis

In order to highlight the structural proteins shared at the inter-specific scale between the isolates of the genus *Betacoronavirus*, thus at the intra-genomic scale of SARS-CoV-2, we have constructed a pan-genome by clustering the sets of proteins encoded in 115 genomes available publicly in NCBI (update: 20-03-2020), including 83 genomes of SARS-Cov-2 and the rest distributed to other species of the same genus. Overall, a total of 1,190 proteins were grouped into a pangenome of 94 orthologous cluster proteins (**Additional file 2: Table S3**), of which ten proteins cluster were shared between SARS-CoV-2 and only three species of the genus *Betacoronavirus* (BatCoV RaTG13, SARS-CoV and Bat Hp-betacoronavirus/ Zhejiang2013). Of these, BatCoV RaTG13 had more orthologous proteins shared with SARS-CoV-2, followed by SARS-CoV with ten and nine orthologous proteins, respectively (**Fig 8A**). It is interesting to note that among all the strains used of *Betacoronavirus*, the ORF8 protein was found in orthology only between SARS-RATG13 and SARS-CoV-2. In addition, the ORF10 protein was found as a singleton for SARS-CoV-2.

**Figure 8:**
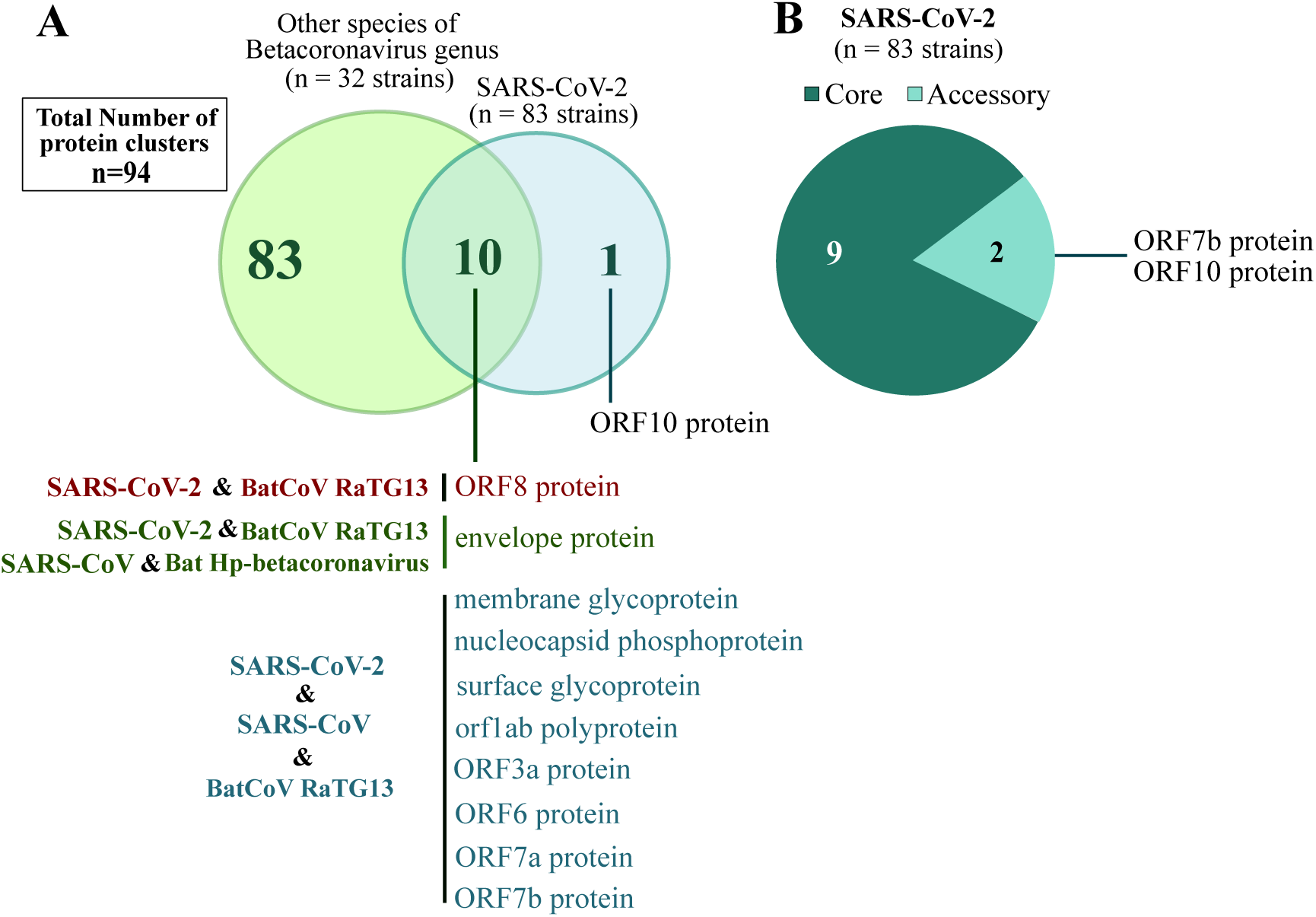
Pangenome analysis of 32 from different Betacoronavirus species and 83 of SARS-CoV-2. (A) The Venn diagram represents the number of core, accessory, and unique proteins inside the Betacoronavirus genus. (B) The pie chart illustrates the core and accessory protein inside the SARS-CoV-2 specie.

On the other hand, the analysis of the pangenome at the intra-genomic scale of 83 isolates of SARS-CoV-2 (**Fig 8B**), showed that ORF7b and ORF10 were two accessory proteins (proteins variable) in SARS-CoV-2 genomes, while the other proteins belonged to the core proteins of SARS-CoV-2 (conserved in all genomes).

## Discussion

The rate of mutations results in viral evolution and variability in the genome, thus allowing viruses to escape host immunity, as well as drugs (20). Initial published data suggests that SARS-CoV-2 is genetically stable (21) which may increase the effectiveness of vaccines under development. The study on the genomic variation of SARS-CoV-2 is very important for the investigation of pathogenesis, disease course, prevention, and treatment of SARS-CoV-2 infection. In this study, we characterized the genetic variations in a large population of SARS-CoV-2 genomes. Our results showed a diversity of mutations detected with different frequencies. Overall, more than 500 non-synonymous mutations in SARS-CoV-2 genomes have been identified. The orf1ab gene having more than half the size of the SARS-CoV-2 genome and is divided into 16 nsp (nsp1-nsp16) (22). We found more than half of recurrent mutations in orf1ab, and a high mutation rate in nsp3, nsp12 and nsp2, with 124, 57 and 46, respectively. Nsp2 and nsp3 were both essential for correcting viral replication errors (23). Thus, recent studies have suggested that mutations falling in the endosome - associated - protein - like domain of the nsp2, could explain why this virus is more contagious than SARS (24).

The replication enzymes nsp12 to nsp16 have been reported as antiviral targets for SARS-CoV (25). In the SARS-CoV-2 genomes, we found that nsp12 to nsp15 harbored nine recurrent non-synonymous mutations. Among them, eight identified as new mutations, including three in nsp12-RNA-dependent RNA polymerase (M4555T, T4847I and T5020I), three in nsp13-Helicase (V5661A, P5703L and M5865V) and two in nsp15-EndoRNAse (I6525T and Ter6668W). However, these new mutations must be taken into account when developing a vaccine using the orf1ab protein sequences as a therapeutic target.

A high number of mutations were identified in the spike protein, an important determinant in pathogenicity that allows the virion attachment to the cell membrane by interacting with the host ACE2 receptor Angiotensin-converting enzyme 2) (26). Among all the frequent mutations in this protein, the V483A mutation has been identified in this receptor and found mainly in SARS-CoV-2 genomes isolated from USA. This result is consistent with the study of Junxian et al. (27). Eight stains from china, USA and France harbored V367F mutation previously described to enhance the affinity with ACE2 receptor (27).

Interestingly, ten hyper variable genomic hotspots with high frequencies of mutated allel detected. Among them, position 11083 (L3606F) detected in NSP6, this protein works with nsp3 and nsp4 by forming double-membrane vesicles and convoluted membranes involved in viral replication (28). Besides, three positions were previously reported by Pachetti et al. (2020), of which the two positions 17858 (M5865V) and 18060 (S5932F) in ORF1ab, and 28881 (R203K) in nucleocapside. Moreover, intraspecies pangenome analysis of SARS-CoV-2 showed that the six of the genes harboring hotspot mutations belong to the core genome.

Thus, under normal circumstances genomic variation increase the viruses spread and pathogenicity. This happens when the virus accumulated mutation enabling its virulence potential (29). Genomic comparison of the studied population allowed us to gain insights into virus mutations occurrence over time and within different geographic areas. In the SARS-CoV virus, the SNP distribution is not random, and it is more dominant in critical genes for the virus (20,30). Our results confirmed what was previously described and elucidate the presence of numerous hotspot mutations. Besides, co-occurrence mutations were also common in different countries all along with singleton mutations. In the case of the China, the singleton mutations are driven by the single group that diverged differently due to the environment, the host, and the number of generations. These mutations are due to the low fidelity of reverse transcriptase (29, 31).

China, US, France and Malaysia contain a high number of specific mutations which may be the cause of a rapid transmission, especially in the US. These specific mutations may also be correlated with the critical condition in US and France.

The clustering of these genomes revealed the spread of clades to diverse geographical regions. We observed an increase of mutations over time following the first dissemination event from China. Specific haplotypes were also predominant to a geographical location, especially in the China. This study opens up new perspectives to determine whether one of these frequent mutations will lead to biological differences and their correlation with different mortality rates.

Among the seven nsp of or1ab hosting sites under selective pressure, only nsp3 and nsp4 contains both residues under positive and negative selection. The modelling of nsp3 domains shows that only the negative selection site 1171 (Thr-353), was located at the conserved macro domain Mac1 (previously X or ADP-ribose 1” phosphatase) (32). This domain has been previously shown to be dispensable for RNA replication in the context of a SARS-CoV replicon (33). However, it could counteract the host’s innate immune response (34). It was proposed that the 3Ecto luminal domain of nsp3 interacts with the large luminal domain of nsp4 (residues 112-164) to induce discrete membrane formations as an important step in the generation of ER viral replication organelles (35, 36). As we have shown previously by the FEL, MEME and FUBAR methods, the orf1ab 2244 site coding for ILE-1426 is under positive selection pressure and since it is located on the luminal 3ecto domain of the nsp3 protein, this can be explained by a possible host influence on the virus in this domain.The results of selective pressure analysis revealed the presence of several negatively selected residues, one of which is located at the receptor-binding domain (GLN-474) and which is known by its interaction with the GLN24 residue of the human ACE2 (Angiotensin-converting enzyme 2) receptor (37). In general, it is well-known that negatively selected sites could indicate a functional constraint and could be useful for drug or vaccine target design, given their conserved nature and therefore less likely to change (38).

## Conclusion

The SARS-CoV-2 pandemic has caused a very large impact on health and economy worldwide. Therefore, understanding genetic diversity and virus evolution become a priority in the fight against the disease. Our results show several molecular facets of the relevance of this virus. We have shown that recurrent mutations are distributed mainly in six SARS-CoV-2 genes with variable mutated allele frequencies. We were able to highlight an increase in mutations accumulation overtime and revealed the existence of three major clades in various geographic regions. Finally, the study allowed us to identify specific haplotypes by geographic location and provides a list of sites under selective pressure that could serve as an interesting avenue for future studies.

## Conflict of interest

The authors declare that they have no competing interests.

## Acknowledgments

We sincerely thank the authors and laboratories around the world who have sequenced and shared the full genome data for SARS-CoV-2 in the GISAID database. All data authors can be contacted directly via www.gisaid.org.

This work was carried out under National Funding from the Moroccan Ministry of Higher Education and Scientific Research (PPR program) to AI. This work was also supported, by a grant to AI from Institute of Cancer Research of the foundation Lalla Salma.

## Notes

http://genoma.ma/covid-19/

### Competing Interest Statement

The authors have declared no competing interest.

